# Genetic dissection of the redundant and divergent functions of histone chaperone paralogs in yeast

**DOI:** 10.1101/586347

**Authors:** Neda Savic, Shawn P. Shortill, Misha Bilenky, David Dilworth, Martin Hirst, Christopher J. Nelson

## Abstract

Gene duplications increase organismal robustness by providing freedom for gene divergence or by increasing gene dosage. The yeast histone chaperones Fpr3 and Fpr4 are paralogs that can assemble nucleosomes *in vitro*, however the genomic locations they target and their functional relationship is poorly understood. We refined the yeast synthetic genetic array (SGA) approach to enable the functional dissection of gene paralogs. Applying this method to Fpr3 and Fpr4 uncovered their redundant and divergent functions: while Fpr3 is uniquely involved in chromosome segregation, Fpr3 and Fpr4 co-operate on some genes and are redundant on others where they impact gene expression and transcriptional processivity. We find that the TRAMP5 RNA exosome is essential in *Δfpr3Δfpr4* yeast and leverage this information to identify Fpr3/4 target loci. Amongst these are the non-transcribed spacers of ribosomal DNA where either paralog is sufficient to establish chromatin that is both transcriptionally silent and refractory to recombination. These data provide evidence that Fpr3 and Fpr4 have shared chromatin-centric functions, especially at nucleolar rDNA. However, their distinct genetic interaction profiles show they also have evolved separate functions outside of the nucleolus.

## Introduction

Gene duplication events play an important role both in driving protein evolution and in providing a mechanism for ensuring the robustness of biological systems. Since the earliest observations of duplications on chromosomes (Darlington & Moffett, 1930; Bridges, 1936) and redundant genes (Kataoka *et al*, 1984; Basson *et al*, 1986), models implicating gene duplication events as complex drivers of evolution have been proposed (Ohno, 1970; Hughes, 1994; Force *et al*, 1999; Francino, 2005; Innan & Kondrashov, 2010). Evolutionary forces can favor the retention of redundant genes for dosage reasons, for example, identical copies of histone and ribosomal genes are present in most eukaryotes. Alternately, duplicated genes provide an opportunity for functional divergence of gene pairs, or paralogs, over time.

*FPR3* and *FPR4* are two *S. cerevisiae* paralogs (Manning-Krieg *et al*, 1994; Shan *et al*, 1994; Benton *et al*, 1994; Dolinski *et al*, 1997) derived from a distant whole genome duplication event (Pemberton, 2006; Wolfe & Shields, 1997; Kellis *et al*, 2004). They code for highly similar proteins (58% identical and 72% similar in amino acid residues) with acidic N-terminal nucleoplasmin-like histone chaperone and C-terminal FK506-binding (FKBP) peptidyl-prolyl isomerase domains (Kuzuhara & Horikoshi, 2004; Xiao *et al*, 2006; Park *et al*, 2014) (Figure 1 A). Both proteins localize to the nucleus and are enriched in the nucleolus (Manning-Krieg *et al*, 1994; Shan *et al*, 1994; Benton *et al*, 1994; Huh *et al*, 2003). Notably, Fpr3 and Fpr4 interact with each other and share some common physical interactors (Krogan *et al*, 2006), including histones (Shan *et al*, 1994; Xiao *et al*, 2006; Nelson *et al*, 2006), and the Nop54 ribosome biogenesis factor (Sydorskyy *et al*, 2005). Additionally, both Fpr3 and Fpr4 are multi-copy suppressors of temperature sensitivity and mating defects resulting from the absence of the Tom1 E3 ubiquitin ligase (Davey *et al*, 2000; Utsugi *et al*, 1999). Therefore, there is good evidence that Fpr3 and Fpr4 operate together and may have redundant functions.

**Figure 1.**
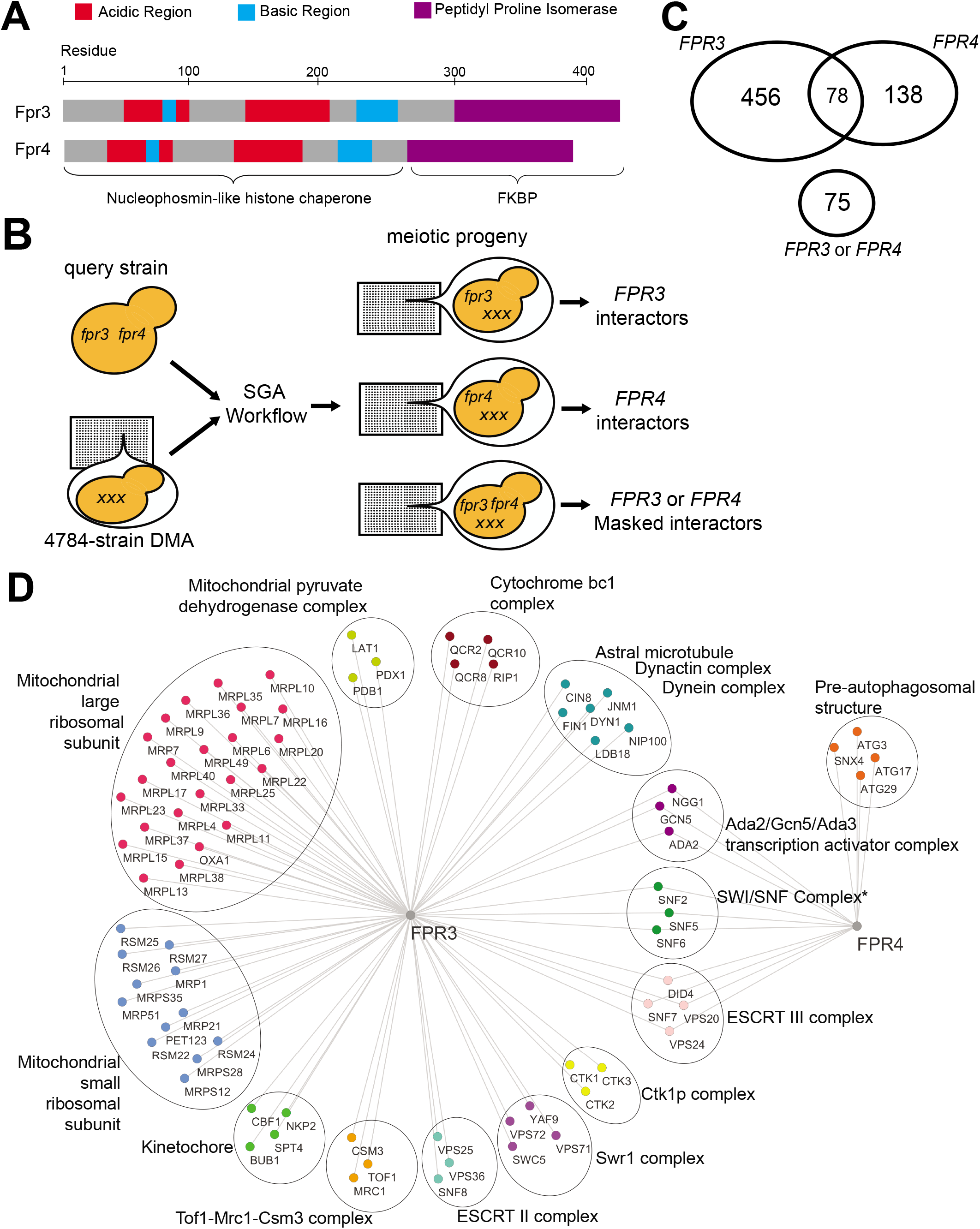
Fpr3 and Fpr4 have separate, co-operative and redundant functions. A. Domain architectures of Fpr3 and Fpr4. Both proteins have an N-terminal nucleoplasmin-like domain with characteristic patches of acidic and basic residues, and a C-terminal peptidyl prolyl isomerase domain. B. Schematic illustrating modified paralog SGA workflow. Spores from a single cross of the double deletion Δ*fpr3*Δ*fpr4* query to the 4784-strain DMA are manipulated to generate three separate sets of meiotic progeny for interactome analysis. C. On top, Venn diagram illustrating numbers of synthetic sick and synthetic lethal genetic interactors unique to *FPR3* and *FPR4*, and shared among both of them. On bottom, number of masked redundant synthetic sick and synthetic lethal genetic interactions only detectable in double deletion Δ*fpr3*Δ*fpr4* mutants. D. Network illustrating complex related ontologies enriched among unique and shared genetic interactors of *FPR3* and *FPR4*. Asterix denotes genetic interactions with the SWI/SNF component coding genes which were confirmed to be shared among Fpr3 and Fpr4 with spotting assays.

There is also evidence that these enzymes are not equivalent. Fpr3 has been identified as a regulator of chromosome dynamics at mitotic and meiotic centromeres. During meiosis, Fpr3 enhances recombination checkpoint delay (Hochwagen *et al*, 2005) and prevents meiotic chromosome synapsis initiation at centromeres (Macqueen & Roeder, 2009). Fpr3 is also required for the degradation of the centromeric histone H3 variant Cse4 (Ohkuni *et al*, 2014). To our knowledge, no reports describe similar data for Fpr4. Taken together, these reports are evidence that Fpr3 and Fpr4 may have functionally diverged.

The comparative impact(s) of Fpr3 and Fpr4 in gene expression are also unclear. While Fpr4 can silence expression of a reporter at ribosomal DNA (rDNA) (Kuzuhara & Horikoshi, 2004) and is involved in transcription induction kinetics through the isomerization of prolines on the amino tails of histones H3 and H4 (Nelson *et al*, 2006), the degree to which Fpr3 regulates transcription has not been described.

In yeast, the loss-of-function phenotypes and genetic interactions of chromatin regulators usually provide insight to their chromatin-centric functions. For example, the yeast histone chaperone coding genes *ASF1* and *RTT106* display clear chromatin-related genetic interactions in synthetic genetic array (SGA) screens (Costanzo *et al*, 2010, 2016). We noted that the genetic interactomes of *FPR3* and *FPR4* contained few chromatin-related hits (Costanzo *et al*, 2010, 2016; Collins *et al*, 2007; Stirling *et al*, 2011; Milliman *et al*, 2012) and hypothesized that the high similarity of these paralogs renders them semi-redundant, masking their genetic interactions.

Here, through a set of comprehensive genetic interaction screens designed for paralogs, we reveal that the functions of Fpr3 and Fpr4 are complex, and include separate, co-operative and redundant functions in chromatin and chromosome biology. Deletion of *Δtrf5*, a key component of the TRAMP5 RNA exosome renders cells reliant on Fpr3/4 for viability, transcriptional processivity and silencing. This strongly suggests that Fpr3/4 and TRAMP5 regulate common RNA transcripts through RNA degradation and chromatin-mediated silencing, respectively. Finally, a major chromatin target for these chaperones is found within the nucleolar rDNA where either protein is sufficient to promote both silencing and genomic stability at the non-transcribed spacer regions. Taken together we have developed a broadly applicable modified SGA approach that can parse out the separate and shared functions of gene paralogs. Applying this to Fpr3/4 has revealed that these histone chaperones have a redundant ancestral function in chromatin regulation of rDNA, however, we also provide evidence that they co-operate and are in the process of functionally diverging.

## Results

### Genetic interactions reveal separate, co-operative, and redundant functions of *FPR3* and *FPR4*

Since *Δfpr3* and *Δfpr4* yeast are viable, but double *Δfpr3Δfpr4* mutants display a synthetic sick phenotype (Costanzo *et al*, 2010; Dolinski *et al*, 1997) we reasoned that their partial redundancy may be masking genetic interactions. To address this and determine the biological processes sensitive to these histone chaperones we performed a modified synthetic genetic array (SGA) screen designed to dissect functional redundancy of gene paralogs (Figure 1 B, see materials and methods). To this end we crossed a dual-query *Δfpr3Δfpr4* double mutant strain to the 4784 strain non-essential yeast deletion mutant array (DMA), so that the fitness of all double (*Δfpr3Δxxx and Δfpr4Δxxx*) and triple (*Δfpr3Δfpr4Δxxx*) mutant meiotic progeny could be measured. The query strain also harbored an episomal *URA3* plasmid with a functional *FPR4* gene to avoid the slow growth phenotype of *Δfpr3Δfpr4* dual deletion, and its vulnerability to suppressor mutations. This plasmid was maintained until the final step of the screen when counter-selection with 5’FOA created the *fpr4* null status. Using standard selection methods, the spores of this single cross were manipulated to generate three separate SGA screens that identified all synthetic lethal/sick interactions with *Δfpr3*, with *Δfpr4* and genes whose disruption only exacerbated fitness of yeast lacking both *Δfpr3Δfpr4*.

We identified 456 and 138 genetic interactors that were unique to either *FPR3* or *FPR4*, respectively, revealing that these paralogs are not equivalent (Figure 1 C top). However, 78 genes interacted with both *FPR3* and *FPR4*, implying that there are specific contexts of paralog co-operativity, that is situations where both histone chaperone is required for function. We also uncovered 75 masked interactors, defined as genes whose deletion only impacts the fitness *Δfpr3Δfpr4* yeast (Figure 1 C bottom). These genes highlight processes when paralog function is redundant. The complete list of these genes and a gene ontology analysis are provided in Appendix file 1 and Appendix file 2 respectively.

*FPR3* genetic interactors include members of the large and small mitochondrial ribosomal subunits (P=3.42×10^−11^ and P=8.38×10^−7^ respectively), the mitochondrial pyruvate dehydrogenase complex (P=6.48×10^−4^), the cytochrome bc1 complex (P=1.49×10^−3^) and components of the ESCRT II endosomal sorting complex (P=8.85×10^−3^) (Figure 1 D). We also identified all three components of the Ctk1 kinase complex (P=1.69×10^−4^), and four components of the Swr1 chromatin remodeler (P=4.45×10^−3^) supporting at least some potential chromatin centric roles of Fpr3. Most notably, we uncovered complexes involved in chromosome segregation such as the astral microtubule (P=2.03×10^−6^), kinetochore (P=2.38×10^−4^), and the Mrc1/Csm3/Tof1 complex (P=1.69×10^−4^) as genetic interactors unique to Fpr3, and not Fpr4. These systems-level data support reports which indicate that Fpr3, but not Fpr4, regulates mitotic and meiotic chromosome dynamics, including those associated with centromeres (Hochwagen *et al*, 2005; Macqueen & Roeder, 2009; Ohkuni *et al*, 2014). Although we identified 138 *FPR4* specific genetic interactions, they fall into limited ontologically related categories. Several genes coding for components of the pre-autophagosome and associated with the process of mitochondrial degradation (P=1.29×10^−3^) were the exception, but the relationship between Fpr4 and these processes is not clear. Taken together the number and nature of the genetic interactions from single query screens suggest that Fpr4 cannot fulfil many of Fpr3’s biological functions, particularly those in chromosome dynamics, and mitochondrial ribosome biology. However, Fpr3 might be competent to substitute for Fpr4 (see below).

Shared genetic interactions would be expected if both paralogs were required for the efficient execution of a biological process. Among genetic interactors common to both *FPR3* and *FPR4* are genes coding for the ESCRT III complex (P=1.44×10^−6^) which functions in endosomal sorting, the Ada2/Gcn5/Ada3 histone acetyltransferase (P=3.57×10^−6^) and the ATP-dependent SWI/SNF chromatin remodeler (Figure 1 D). Shared genetic interactions with the SWI/SNF remodeler were confirmed using spotting assays (data not shown). The proposed co-operation of Fpr3 and Fpr4 is supported by the fact these proteins co-purify (Krogan *et al*, 2006) and, like nucleoplasmin, have the intrinsic propensity to form oligomers (Edlich-Muth *et al*, 2015; Dutta *et al*, 2001; Koztowska *et al*, 2017). Thus, these shared genetic interactions with known chromatin regulatory complexes support published protein complex data and indicate that Fpr3 and Fpr4 likely co-operate through heteromeric complexes in some contexts.

75 masked genetic interactions are only detectible in double *Δfpr3Δfpr4* mutants (Figure 1 C bottom). These genes are essential only when both paralogs are absent, and thus highlight processes in which Fpr3 and Fpr4 are redundant. Most notably, these interactors include three non-essential components of the TRAMP5 nuclear RNA exosome (*TRF5*, *AIR1*, and *RRP6*) (Figure 2 A), an RNA surveillance factor that recognizes, polyadenylates and degrades aberrant RNA transcripts (Figure 2 B) (LaCava *et al*, 2005; San Paolo *et al*, 2009; Houseley & Tollervey, 2008; Wery *et al*, 2009). We independently confirmed synthetic sickness of *Δfpr3Δfpr4* with *Δrrp6* and *Δtrf5* using growth curves (Figure 2 C). This demonstrates that Fpr3 and Fpr4 have redundant biological functions likely involving the negative regulation of RNAs.

**Figure 2.**
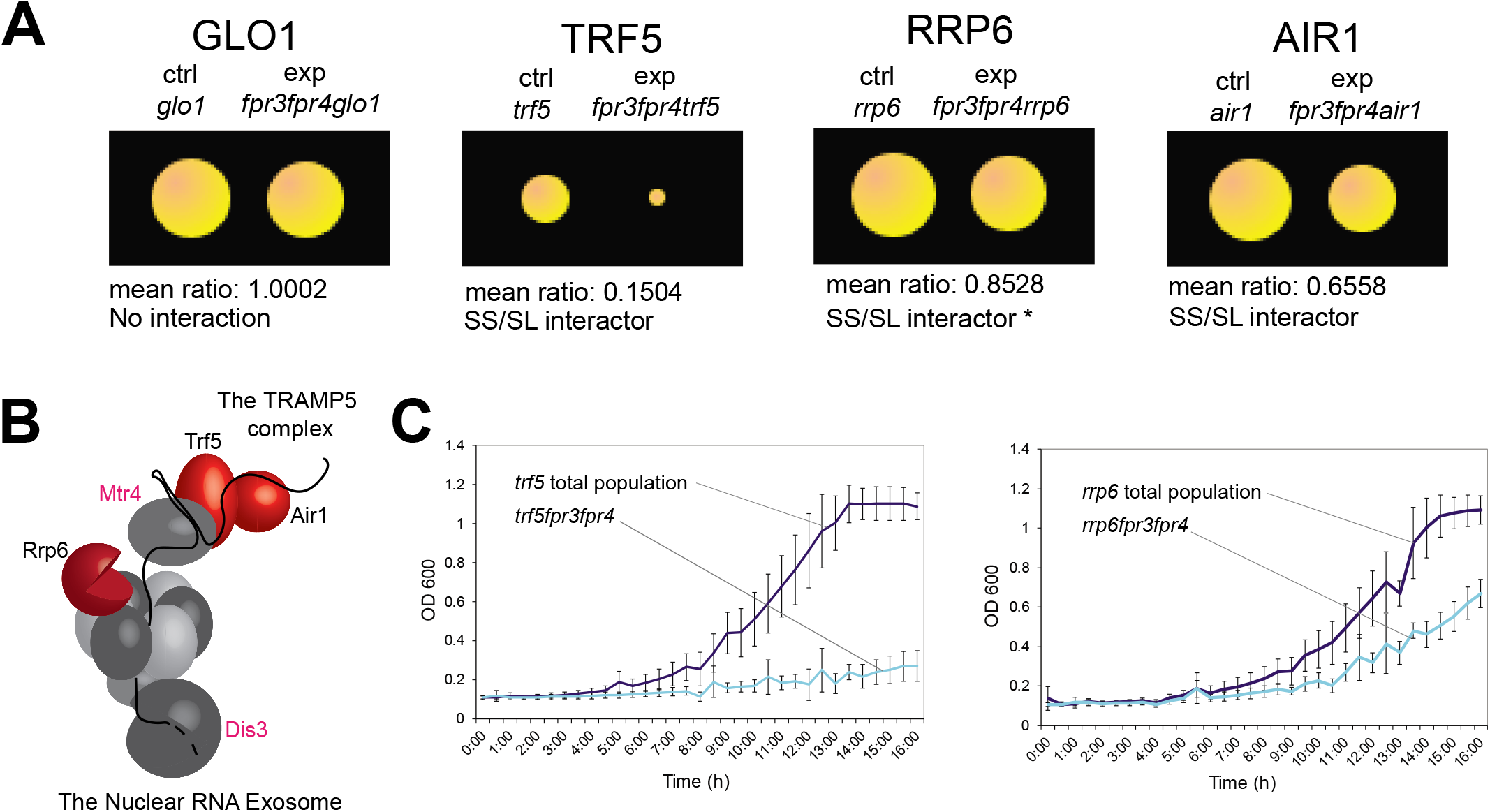
The TRAMP5 nuclear RNA exosome is a masked genetic interactor of *FPR3* and *FPR4*. A. Mean colony size ratios of experimental (Δ*fpr3*Δ*fpr4*Δ*xxx*) triple mutants relative to control Δ*xxx* total haploid meiotic progeny for select redundant synthetic sick or lethal genetic interactors. Asterix indicates that 2/3 replicates for the Δ*fpr3*Δ*fpr4*Δ*rrp6* deletion mutant were below the synthetic sick/ lethal cut-off threshold. B. Illustration of the TRAMP5 complex (top right) interacting with the nuclear RNA exosome (bottom left). Complex components coded for by redundant genetic interactors of *FPR3* or *FPR4* are colored red. Pink text labels indicate components of complex coded for by essential genes. Illustration is adapted from (Wolin *et al*, 2012). C. Growth curves depicting OD600 vs time for select triple deletion mutants and corresponding total haploid meiotic progeny control populations.

### Suppressor genetic interactions of *FPR3* and *FPR4*

The SWI/SNF and Ada2/Gcn5/Ada3 complexes are particularly important for the fitness of *Δfpr3* and *Δfpr4* yeast (Figure 1 D). In support of a chromatin defect underlying these phenotypes, we found that several genetic suppressors (Figure 3), which alleviate the slow growth phenotype of *Δfpr3Δfpr4* yeast, are themselves chromatin modifiers including: three NAD+ dependent histone deacetylases (P= 6.33×10^−5^), Hos2, Hda1 and Hos3; three of the four components of the HIR replication-independent nucleosome assembly complex (P=1.29×10^−5^), Hir1, Hpc2, and Hir3; and Swd3 and Sdc1, two of the eight components of the Set1/COMPASS histone H3K4 methylase complex, (P= 5.87×10^−3^). We note that the Swd2 subunit of COMPASS is encoded by an essential gene and the *Δset1* knockout is not present in our deletion strain collection. It is particularly notable that we find histone deacetylases enriched among suppressor interactions and histone acetyltransferases among synthetic sick and lethal interactions. The presence of both aggravating and alleviating chromatin-related genetic interactions in our modified SGA screen is consistent with a chromatin-centric mode of action for Fpr3 and Fpr4.

**Figure 3.**
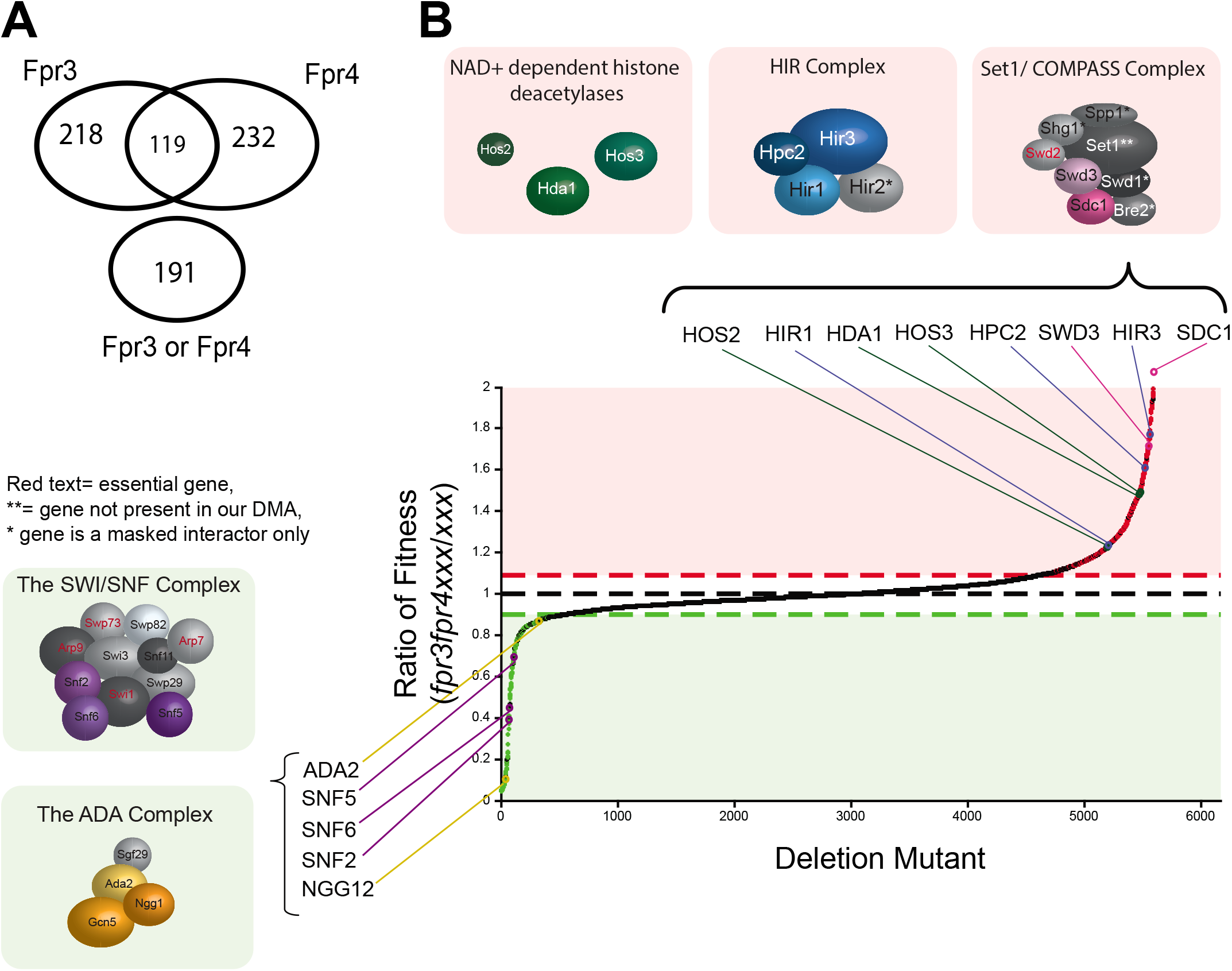
Suppressor genetic interactions support chromatin-centric functions for Fpr3 and Fpr4. A. On top, Venn diagram illustrating numbers of suppressor interactors unique to *FPR3* and *FPR4* and shared among both of them. On bottom, number of masked redundant suppressor genetic interactions only detectable in double deletion Δ*fpr3*Δ*fpr4* mutants. B. Plot of fitness ratios for all Δ*fpr3*Δ*fpr4*Δ*xxx* triple mutants relative to Δ*xxx* total haploid meiotic progeny controls. Green dots indicate all synthetic sick/ lethal genetic interactions, red dots indicate all suppressor genetic interactions. Threshold cut-offs are indicated by red and green dashed horizontal lines. Fitness ratios associated with genes coding for components of chromatin modifiers are labeled and accompanied with schematic illustrations of complex components coded for by the synthetic sick genetic interactors (illustrated in green boxes) and suppressor genetic interactors (illustrated in red boxes). Components coded for by interacting genes are colored. Components coded for by non-interacting genes are black and white. Red text illustrates components coded for by essential genes absent from the non-essential yeast DMA.

### Fpr3 and Fpr4 have shared and separate transcriptional targets

The genetic interactions of Fpr3 and Fpr4 with known chromatin modifiers suggest that they regulate transcription. Consistent with this, Fpr4 directly represses transcription from a reporter gene both in an artificial recruitment assay (Park *et al*, 2014) and integrated in the rDNA repeats of yeast (Kuzuhara & Horikoshi, 2004). Fpr4 is also bound to multiple genomic locations (Nelson *et al*, 2006; Kuzuhara & Horikoshi, 2004). To determine the impact of these proteins on transcription genome-wide, we sequenced the ribo-minus fraction of RNAs from wt, *Δfpr3, Δfpr4* and *Δfpr3Δfpr4* yeast (Appendix file 3, and Figure 4 A). We included a *Δsir2* strain as a control, which in our analysis displays 854 differentially expressed genes using a lenient cut-off of 1.3 fold. This number and nature of Sir2-regulated genes is in good agreement with previous reports of Sir2 regulated genes and binding sites (Ellahi *et al*, 2015; Li *et al*, 2013).

**Figure 4.**
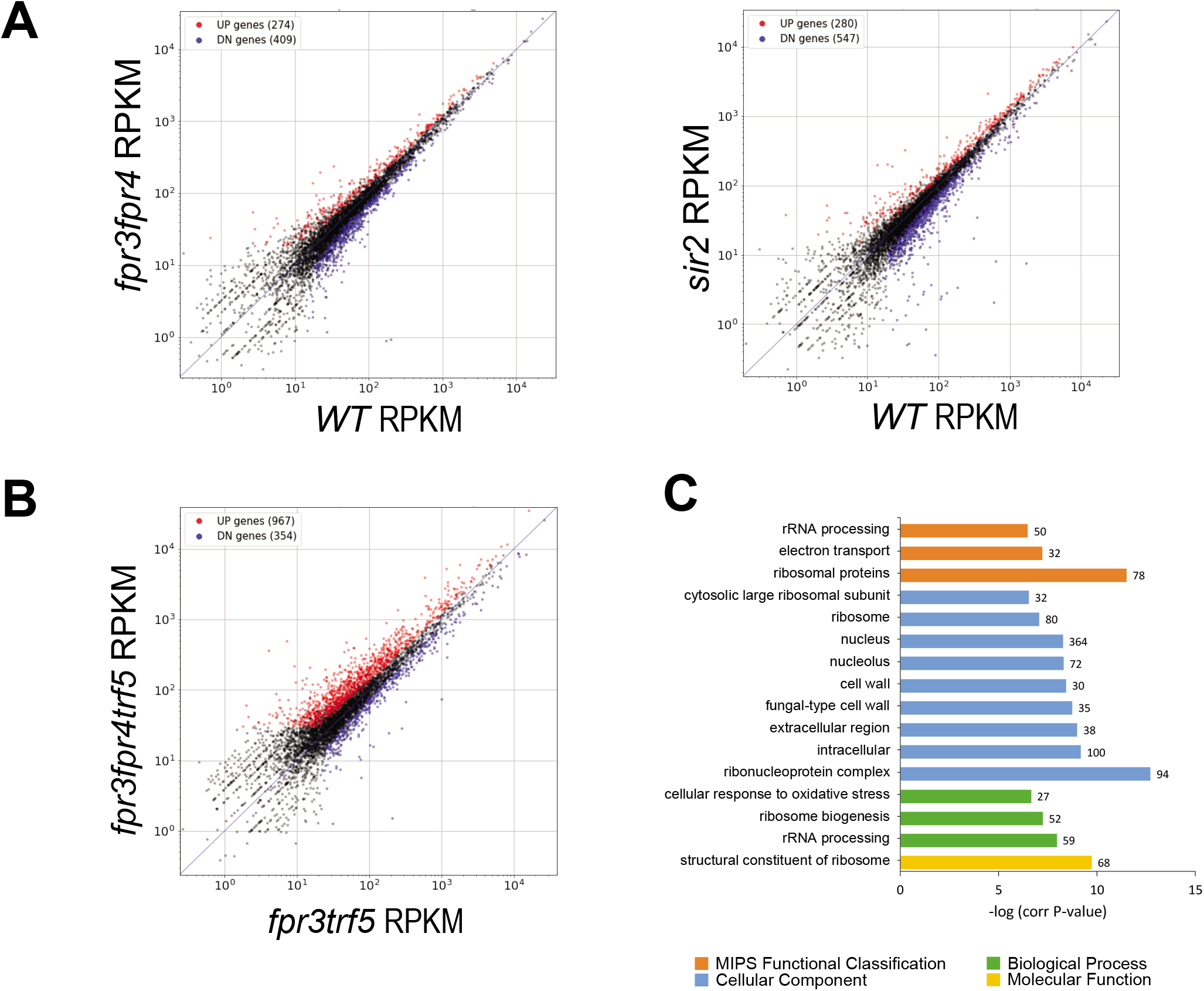
Fpr3 and Fpr4 negatively regulate ribosomal protein and rRNA processing genes. A. Scatter plots indicating the correlation of gene expression between wt and Δ*fpr3*Δ*fpr4* and wt and Δ*sir2* deletion mutants. B. Scatter plots indicating the correlation of gene expression between Δ*fpr3*Δ*trf5* double mutants and Δ*fpr3* Δ*fpr4*Δ*trf5* triple deletion mutants. C. Gene ontology enrichment analysis for upregulated transcripts in Δ*fpr3* Δ*fpr4* Δ*trf5* triple deletion mutants. Enriched genes were classified by molecular function, biological process, cellular component, and MIPS functional database classification by FunSpec (http://funspec.med.utoronto.ca/).

Single deletion mutants of *Δfpr3* and *Δfpr4* had 524 and 549 differentially expressed genes, respectively (Appendix file 3). Surprisingly, double *Δfpr3Δfpr4* mutants did not exhibit a major additive effect with only 683 differentially regulated genes. In each of the three above experiments, approximately 1/3 of differentially expressed genes were upregulated. These genes represent transcripts repressed by the histone chaperone(s) and include members of the cytosolic large ribosomal subunit (P=3.40×10^−11^ in *Δfpr3* mutants, P=8.94×10^−8^ in *Δfpr4* mutants, and P=4.41×10^−12^ in *Δfpr3Δfpr4* mutants), components of the cytosolic small ribosomal subunit (P=8.99×10^−6^ in *Δfpr3* mutants, P=5.84×10^−6^ in *Δfpr4* mutants, and 2.69×10^−10^ in *Δfpr3Δfpr4* mutants) and components of the fungal-type cell wall (P=1.47×10^−4^ in *Δfpr3* mutants, P=4.90×10^−4^ in *Δfpr4* mutants, and P=2.56×10^−3^ in *Δfpr3Δfpr4* mutants). Some of the most differentially expressed genes (up to 60 fold) include proteins involved in phosphate metabolic processes such as the PHO5 and PHO11/12 acid phosphatases, and the phosphate transporters PHO89, PHO84 and PIC2 (P=9.77×10^−7^ in *Δfpr3* mutants, P=3.51×10^−5^ in *Δfpr4* mutants, and P=1.77×10^−4^ in *Δfpr3Δfpr4* mutants, Appendix file 4).

Two-thirds of differentially regulated genes are positively regulated by Fpr3/4. These include fungal type cell wall organization factors (P=1.36×10^−5^ in *Δfpr3* mutants, P=2.14×10^−3^ in *Δfpr4* mutants, and P=1.62×10^−3^ in *Δfpr3Δfpr4* mutants); proteins involved in iron ion homeostasis (P=1.91×10^−7^ in *Δfpr3* mutants, P=2.53×10^−5^ in *Δfpr4* mutants, and P=9.86×10^−5^ in *Δfpr3Δfpr4* mutants); and pheromone response, mating type determination and sex specific proteins (P=3.26×10^−8^ in *Δfpr3* mutants, P=2.42×10^−4^ in *Δfpr4* mutants, and P=2.20×10^−5^ in *Δfpr3Δfpr4* mutants).

Since roughly one third of Fpr3 regulated transcripts are also regulated by Fpr4, and vice versa, we conclude that, on these genes, transcriptional regulation requires both Fpr3 and Fpr4. In other words, Fpr3 and Fpr4 co-operate in these contexts. Like the SWI/SNF complex, the impact of the Fpr3 and Fpr4 histone chaperones can be positive and negative, depending on the gene.

### The TRAMP5 RNA exosome masks the impact of Fpr3/4

Considering that the TRAMP5 nuclear RNA exosome is essential in yeast lacking Fpr3 and Fpr4, we wondered whether this RNase could be masking changes in the *Δfpr3Δfpr4* transcriptome. To test this idea, we sequenced RNA from isogenic *Δtrf5* deficient yeast from our SGA screen, comparing those with a functional *Fpr4(Δfpr3Δtrf5*) to those with neither Fpr3/4 protein (*Δfpr3Δfpr4Δtrf5*). This analysis, designed to reveal Fpr4 regulated RNAs, uncovered a total of 1321 differentially expressed genes (967 upregulated and 354 downregulated) (Figure 4 B). The increase in upregulated transcripts in this experiment supports the hypothesis that Fpr4 negatively regulates a breadth of genes, and that these RNAs are also substrates for the TRAMP5 RNA exosome. As expected, upregulated genes coding for protein components of the cytosolic ribosome (P=3.21×10^−12^) (including the cytosolic large ribosomal subunit P=3.00×10^−7^ and the cytosolic small ribosomal subunit P=9.48×10^−4^) and genes associated with rRNA processing (P=1.14×10^−8^) are highly enriched as Fpr4 targets. Also enriched were genes coding for constituents of the fungal-type cell wall (P=1.87×10^−4^) and the electron transport chain (6.12×10^−8^) (Figure 4 C). Taken together the ontologies associated with upregulated transcripts in *Δfpr3Δfpr4Δtrf5* triple mutants indicate that Fpr3 and Fpr4 negatively regulate discreet subsets of genes, particularly those involved in ribosome biogenesis. That Fpr3/4 and TRAMP5 negatively regulate overlapping transcripts provides a potential explanation for their synthetic lethality.

### A signature of abortive transcription in *Δfpr3Δfpr4* yeast

Further interrogation of the transcriptome data reveals evidence for Fpr3 and Fpr4 in transcriptional processivity: approximately 40% of differentially expressed genes in *Δfpr3Δfpr4Δtrf5* yeast displayed an accumulation of reads towards the 5’ end of the annotated transcript. Subsequent bioinformatic analysis of the total transcriptomes of *Δfpr3Δfpr4Δtrf5* and *Δfpr3Δtrf5* mutants revealed that this 5’-biased asymmetry is widespread, and detectable in genes irrespective of their net change in transcription (Figure 5 A). Two example genes illustrating this asymmetry signature are presented in Figure 5 B; *SSF1* codes for a constituent of the 66S pre-ribosome and is required for large ribosomal subunit maturation, while *UTP1* codes for a component required for proper endonucleolytic cleavage of 35S rRNA. The paired-end tag coverage associated with both of these genes, but not the *IDP1* gene (Figure 5 C), displays the characteristic 5’assymetry in *Δfpr3Δfpr4Δtrf5* yeast. This signature demonstrates Fpr3 and Fpr4 negatively regulate transcription from many promoters and suggests that in the absence of these histone chaperones, transcription can initiate, but may not proceed to completion. That these abortive mRNAs are only readily detectable in the absence of Trf5 indicates that RNA exosomes can mask subtle transcriptional defects (Figure 5 D).

**Figure 5.**
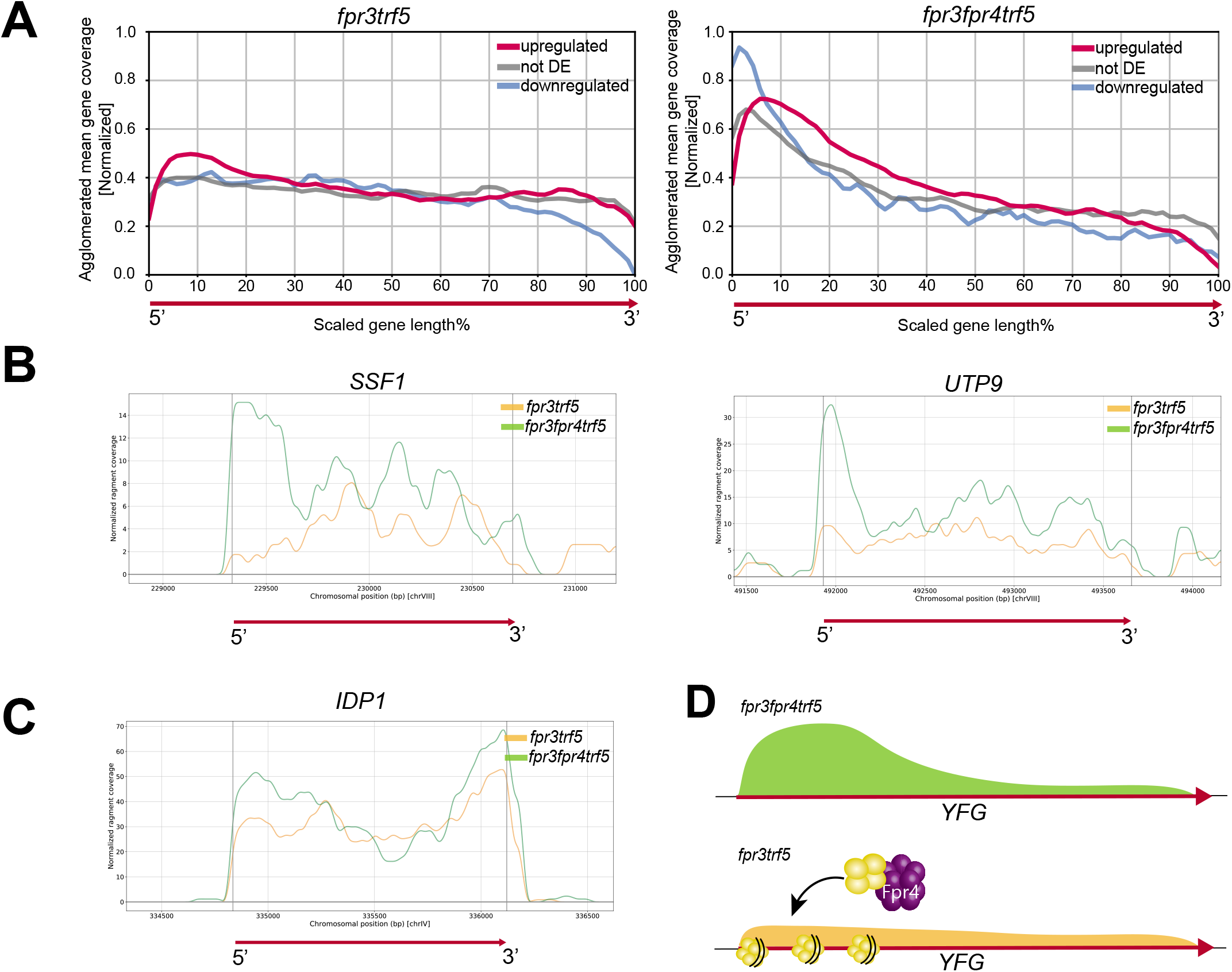
A signature of abortive transcription is present in Δ*fpr3*Δ*fpr4* yeast. A. Plot of total averaged upregulated, downregulated and unchanged transcripts generated form Δ*fpr3*Δ*trf5* double mutants (left) and Δ*fpr3* Δ*fpr4* Δ*trf5* triple mutants (right). B. Read maps illustrating two examples of genes showing a signature of abortive transcription: *SSF1* (left), *UTP9* (right). C. Read map illustrating an example of a non-differentially expressed gene without a signature of abortive transcription *IDP1*. D. Model illustrating Fpr4 building chromatin at gene promoters.

### Fpr3 and Fpr4 inhibit transcription from the non-transcribed spacers of ribosomal DNA

The ribosomal DNA locus in yeast consists of a series of 150-200 tandem repeats of a 9.1kb unit containing the 35S and the 5S rRNAs each separated by two non-transcribed spacer sequences (NTS1 and NTS2) (Figure 6 A top) (Johnston *et al*, 1997). Given the nucleolar enrichment of Fpr3 and Fpr4, and the ability of Fpr4 to repress reporter expression from rDNA (Kuzuhara & Horikoshi, 2004), we asked if yeast lacking Fpr3 and Fpr4 also display transcriptional defects at rDNA. While our RNA-seq analysis was performed on ribo-minus RNA, reads from rRNA are readily detected (presumably from incomplete rRNA depletion) and indicated no change in rRNA production, which we have also observed in Northern and qRT-PCR analyses (data not shown). However, disruption of both Fpr3 and Fpr4 has a profound impact on silencing at NTS1 and NTS2 (Figure 6 A). Transcripts from both strands of NTS1 and NTS2 accumulate in *Δfpr3 Δfpr4 Δtrf5* strains. Consistent with previous reports (Kuzuhara & Horikoshi, 2004), we also find that the repression of a URA3 reporter gene integrated at the NTS1 region of rDNA requires Fpr3 and Fpr4 (Figure 6 B). Taken together, these results support a model where Fpr3 and Fpr4 establish a transcriptionally silent chromatin structure at rDNA.

**Figure 6.**
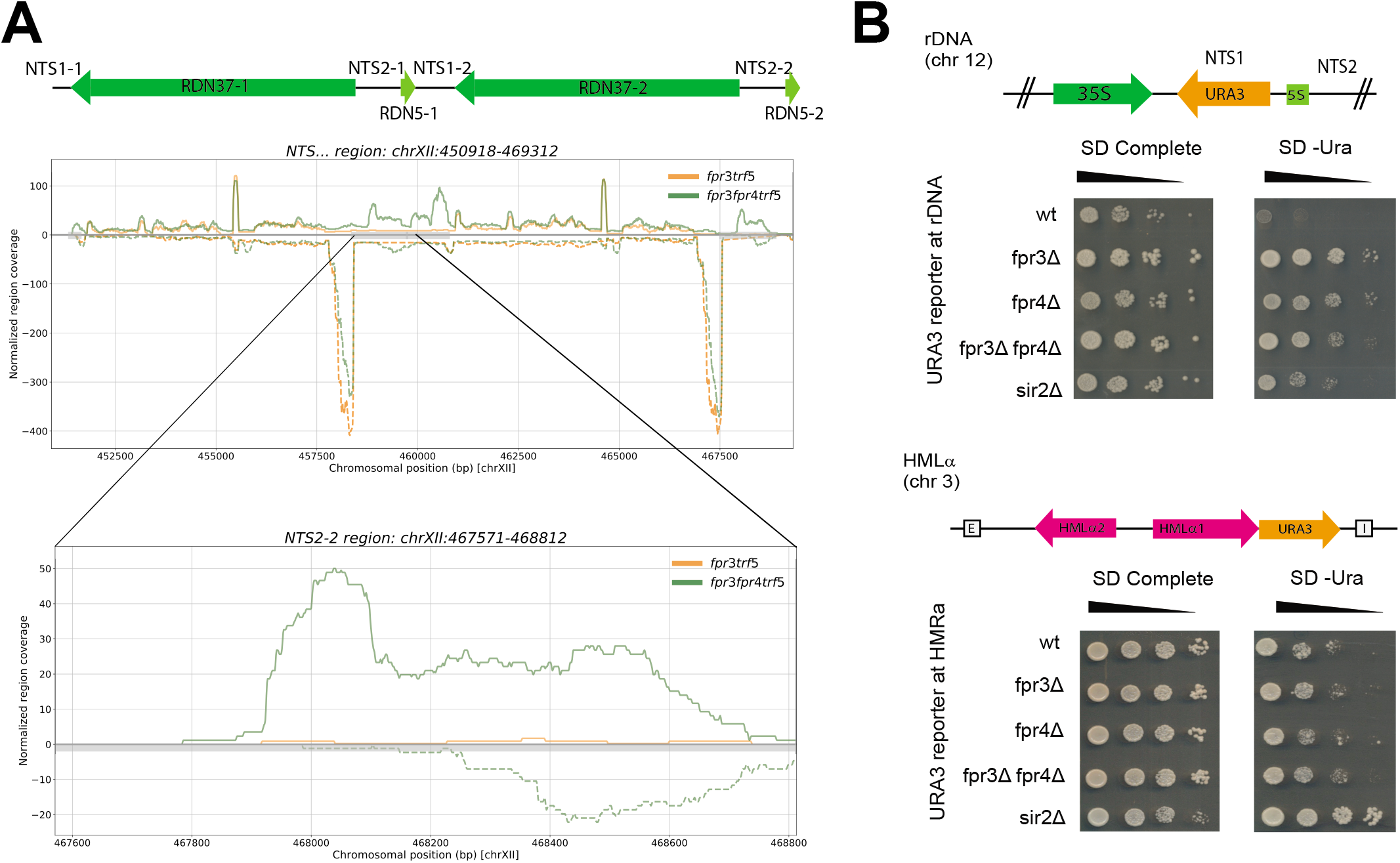
Fpr3 and Fpr4 silence the non-transcribed spacers (NTS) of rDNA. A. Read maps illustrating transcripts generated from both strands of one of the tandem rDNA repeats in <u>Δ*fpr3*Δ*trf5*</u> and Δ*fpr3*Δ*fpr4*Δ*trf5* cells. Transcripts generated from the *NTS2* locus are prezented in the zoomed-in panel. B. Ten-fold serial dilution spotting assays of single and double gene deletion mutants in strain backgrounds carrying a *URA3* reporter integrated either within *NTS1* spacer of rDNA or at the *HMRa* locus. Plates were grown on either standard defined complete media or on standard defined media lacking uracil for 2 days at 37°C.

### Fpr3 and Fpr4 contribute to genomic stability at ribosomal DNA

Ribosomal RNAs comprise approximately 80% of the total RNA in yeast; accordingly the ~ 50% of rDNA tandem repeats that are transcribed in a given cell are the most heavily transcribed, and nucleosome-free, genes in the cell (Nomura *et al*, 2004; Warner, 1999; Vogelauer *et al*, 2000). Reciprocally, the adjacent rDNA non-transcribed spacers (NTS) and inactive rDNA repeats are chromatinized and potently silenced. This arrangement is thought to generate a chromatin template that is refractory to recombination between rDNA repeats and the deleterious loss of rDNAs from chromosome 12, which is a major driver of yeast replicative aging (Sinclair & Guarente, 1997). For this reason, failure to generate heterochromatin environments at rDNA, as occurs in *Δsir2* histone deacetylase mutants, decreases genomic stability at this locus (Gottlieb & Esposito, 1989; Kobayashi *et al*, 2004). We reasoned that if Fpr3 or Fpr4 were silencing the NTS regions via a mechanism that involves chromatin structure, that yeast lacking these enzymes should also exhibit genomic instability at this locus. To test this hypothesis, we introduced *Δfpr3Δfpr4 and Δsir2* deletions into a strain with a reporter gene (URA3) integrated at NTS1 (Van Leeuwen *et al*, 2002; van Leeuwen & Gottschling, 2002). First, URA+ status of each strain was confirmed, followed by growth in non-selective media (YPD) for two days to permit reporter loss. Next, *ura-* cells were isolated on 5’FOA and ~96 colonies picked using a colony picking robot. These *ura*-cells could arise in two ways: epigenetic silencing of *URA3* at NTS1, or from URA3 gene loss via recombination (Figure 7 A top). To discriminate between these events, we replica plated these individual isolates to media lacking uracil, where growth would indicate that the *URA3* phenotype was a consequence of epigenetic silencing. Reciprocally, isolates that failed to grow would represent reporter loss events (Figure 7 A). These propagation assays revealed that normally, the rate of epigenetic switching of URA3 is much higher than reporter loss: 82% of *ura*- isolates still have a *URA3* gene at the end of our propagation assay as exemplified growth in the absence of uracil (Figure 7 B and C), and by PCR of genomic DNA (not shown). As expected, *Δsir2* yeast are unable to establish silent chromatin at NTS1, and can only grow on 5’FOA via loss of the reporter. Finally, we observe that *Δfpr3Δfpr4* yeast are compromised in their ability to silence *URA3* epigenetically: only 30% of 5’FOA resistant colonies retain the *URA3* gene. Thus, in *Δfpr3Δfpr4* yeast recombination and URA3 reporter gene loss are more frequent than epigenetic silencing. This observation supports a model where Fpr3 and Fpr4 build chromatin structures at the NTS regions of rDNA locus. These structures are critical to maintaining genome stability at rDNA.

**Figure 7.**
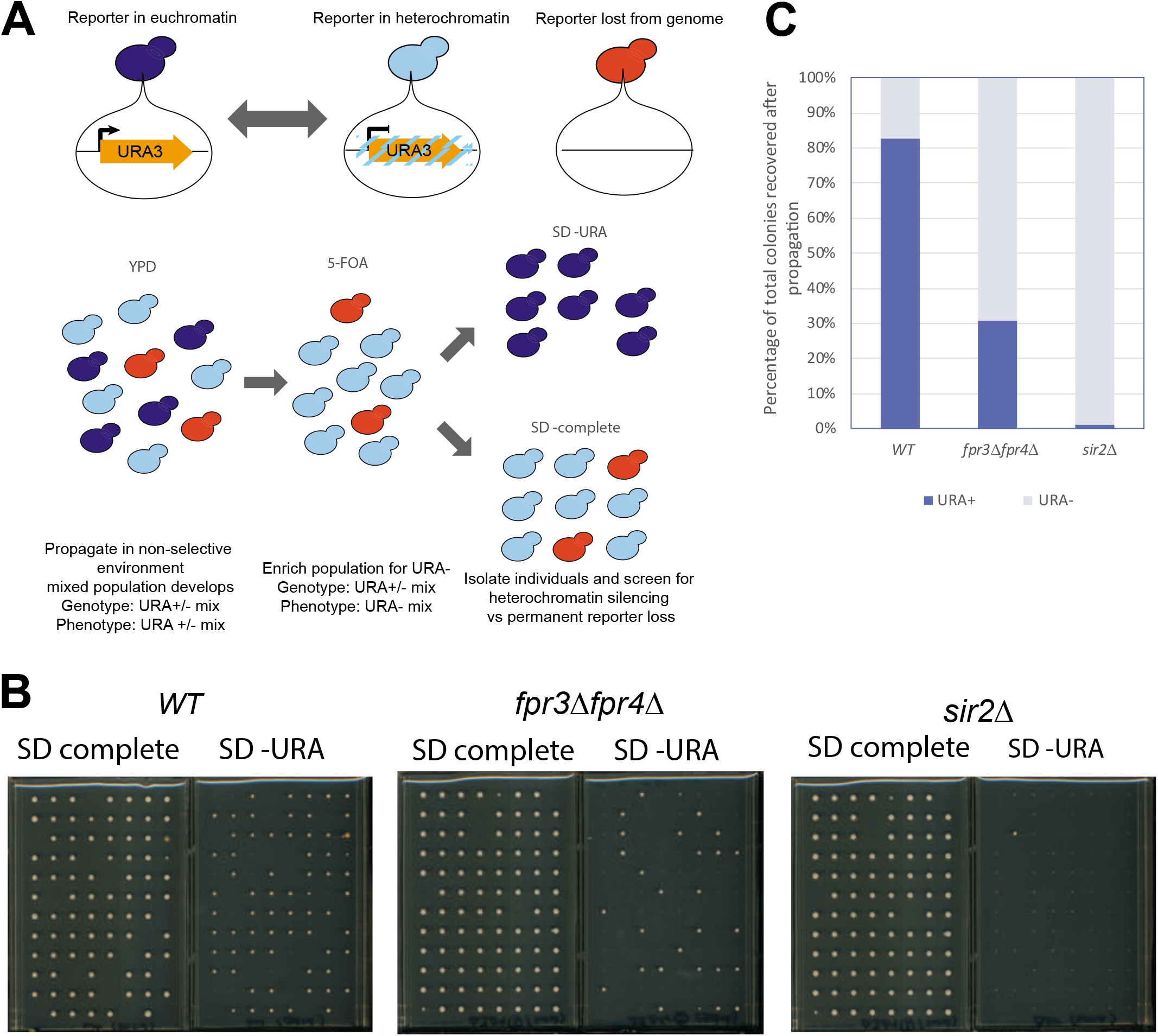
Fpr3 and Fpr4 are required for genomic stability at the rDNA locus. A. Diagrams illustrating the propagation experiment carried out to assess frequency of reporter loss. In a given population of cells, under non-selective conditions, *URA3* may be in an accessible euchromatin-like environment and therefore expressed (dark blue cells), in an inaccessible heterochromatin-like environment and therefore silenced (light blue cells), or it may have been permanently lost from the genome via recombination between repeats (orange cells). B. Images of the 96 individuals selected for after propagation on SD-complete control media and on SD-URA experimental media. Those growing on the experimental media represent the fraction of the population in which the reporter was epigenetically silenced. Those that fail to grow indicate permanent loss of the reporter. C. Percentage of total colonies recovered after strain propagation that have retained or lost the ability to grow on SD-complete media.

## Discussion

Gene duplication events play a critical role in protein and organism evolution. However, the high similarity of duplicated genes can lead to complete or partial compensation when one paralog is deleted, as is in the case in conventional genetic interaction analysis. Here we present a dual-query SGA screening approach where one genetic cross can report the separate and redundant genetic interactions of each paralog. Using this approach on two nucleoplasmin-like histone chaperones revealed that they perform separate, cooperative, and redundant chromatin-related functions. Given that approximately 13% of yeast protein coding genes are duplicates (Wolfe & Shields, 1997), this approach may have wide applications to other studies of yeast paralogs.

The genetic interactions annotated here support a unique function for Fpr3 in orchestrating centromeric chromatin dynamics during chromosome segregation. This is fully consistent with existing literature (Hochwagen *et al*, 2005; Macqueen & Roeder, 2009; Ghosh & Cannon, 2013; Krogan *et al*, 2006; Ohkuni *et al*, 2014). Our comparative analyses provide additional systems-level evidence that this role is not shared with Fpr4 indicting that Fpr3, potentially as homo-oligomers, may regulate chromatin in a way that impacts chromosome segregation (Macqueen & Roeder, 2009; Hochwagen *et al*, 2005). Furthermore, the fact that *Δfpr3Δfpr4* double mutants display fewer genetic interactions than single gene *Δfpr3* mutants (Appendix file 1) indicates that Fpr4 may be toxic in the absence of Fpr3 (Ohkuni *et al*, 2014). This model predicts that, in the absence of Fpr3, the partial engagement or modification of chromatin by Fpr4 is deleterious.

SWI/SNF complex members are shared interactors of Fpr3 and Fpr4, appearing as hits in both single mutant screens. These results could be explained by reduced dosage of a histone chaperone. Alternately, these genetic interactions are consistent with a model where Fpr3 and Fpr4 act together to chaperone nucleosomes, facilitating chromatin dynamics as SWI/SNF does. Whether this means that the chaperones operate together in a sequence of events, such as the removal and subsequent redeposition of nucleosomes during transcription or, in concert as a hetero-oligomeric complex, is not yet clear. The fact that Fpr3 and Fpr4 co-purify (Krogan *et al*, 2006) supports the latter model, but does not exclude the former.

The repression of several PHO genes in rich media requires both Fpr3 and Fpr4. The PHO5, PHO11/12 acid phosphatases, and the PHO89, PHO84 and PIC2 phosphate transporters are intimately linked to the metabolism of both phosphate and intracellular polyphosphate stores. It is therefore intriguing that both Fpr3 and Fpr4 were recently identified as two of the major polyphosphorylated proteins in yeast along with several proteins in an evolutionarily conserved network of ribosome biogenesis factors (Bentley-DeSousa *et al*, 2018). The precise sites of Fpr3/4 polyphosphorylation and their impact on function is not yet clear. However, the identification of the well-studied *PHO5* gene as an Fpr3 and Fpr4 target provides an ideal system for determining the impact of this new post-translational modification on these histone chaperones.

The yeast TRAMP5 complex recognizes and polyadenylates aberrant RNA transcripts in order to target them for degradation by the Rrp6 ribonuclease (Karyn Schmidt and J. Scott Butler, 2013). TRAMP5 targets include both ribosomal protein coding mRNAs and cryptic unstable transcripts generated from intragenic sites on the genome including those within the ribosomal DNA locus (Reis & Campbell, 2007; San Paolo *et al*, 2009; Wery *et al*, 2009; LaCava *et al*, 2005). Here we found that deletion of *Δtrf5* enabled the detection of an unexpected transcriptome signature in *Δfpr3Δfpr4* yeast where there is a bias in RNA-seq reads towards the 5’ end of genes. This means that Fpr3/4 redundantly promote the transcriptional elongation process. It is noteworthy that these reads appear to cover the first 1-3 nucleosomes of genes because Fpr3/4 have evolved basic surface features to permit nucleosome binding (Leung *et al*, 2017) and that Fpr4 was previously shown to be important for the kinetics of transcriptional induction (Nelson *et al*, 2006). Thus, the nucleosomes immediately downstream of the transcriptional start site are candidates targets of Fpr3/4. This regulation could involve either depositing histones within promoters to inhibit transcriptional initiation or the eviction of nucleosomes from sequences downstream of the promoter in order to remove nucleosome blocks to the polymerase. These models are currently under investigation.

Fpr3/4 have the greatest impact on the steady-state levels of mRNAs encoding ribosomal protein genes and rRNA processing machinery. Thus, Fpr3/4 may function as master regulators of ribosome biogenesis by coordinating both ribosomal protein abundance and rRNA processing. Given that many ribosomal and rRNA processing protein genes are driven by common regulators, Fpr3/4 may recognize common DNA sequences or transcription factors to accomplish this function (Fermi *et al*, 2016). It appears that at least some elements of this ribosomal biogenesis function are conserved in the human nuclear FKBP25 protein (Gudavicius *et al*, 2014; Dilworth *et al*, 2017).

In addition to regulating the transcription of protein coding genes Fpr3 and Fpr4 restrict transcription from the non-transcribed spacers (NTS) sequences of ribosomal DNA. This is consistent with both their nucleolar enrichment and data indicating that they inhibit transcription of exogenous reporters at NTS2 in yeast (Kuzuhara & Horikoshi, 2004) and endogenous 18S rDNA in plants (Li & Luan, 2010). In yeast the NTS loci contain important DNA sequence features including as two terminators for the RNA PolI transcribed RDN35 repeat, a replication fork barrier site, and an autonomous replication site. Two separate observations suggest that Fpr3 and Fpr4 function redundantly to build chromatin at rDNA in order to insulate DNA at these spacers. First, yeast lacking both paralogs accumulate large amounts of aberrant NTS RNA transcripts, and these RNAs are templated by both DNA strands. Second, consistent with a chromatin structure defect underpinning this phenomenon, we find that the rDNA locus in *Δfpr3Δfpr4* yeast is also hyper-recombinogenic (Figure 6). Thus, Fpr3 and Fpr4 are histone chaperones of particular importance at the 100-200 rRNA repeats where they mediate the stability and silencing of spacers between the most heavily transcribed sequences in the cell. How these chaperones regulate chromatin structure at this locus, and how the structure differs from other targets in the nuclear genome remain open questions.

## Materials & Methods

### Yeast strains and plasmids

Yeast strains used in this study are described in Appendix file 5. Strains in the MAT a non-essential yeast deletion collection (DMA) used for the SGA analysis are all isogenic to BY4741 and were purchased from Thermofisher Dharmacon. The plasmid rescued double genomic deletion Δ*fpr3*Δ*fpr4* SGA query strain (YNS 35) was created in a Y7092 genetic background as follows. The endogenous *FPR4* locus on a Y7092 wt strain was replaced with a nourseothricin resistance (*MX4-NATR*) PCR product deletion module. The resulting single gene Δ*fpr4* deletion mutant was subsequently transformed with prs316 FPR4: a single copy *URA3* marked shuttle vector carrying an untagged full length copy of the *FPR4* open reading frame with endogenous promoter and terminator (originally described in (Nelson *et al*, 2006)). The endogenous *FPR3* locus on this plasmid rescued Δ*fpr4* deletion mutant was subsequently replaced with a *LEU2* PCR product deletion module.

Triple deletion mutants: Δ*rrp6*Δ*fpr3*Δ*fpr4* and Δ*trf5*Δ*fpr3*Δ*fpr4* and their corresponding mixed population total haploid meiotic progeny controls used in the validating growth curves were generated from the SGA cross (see below).

Single gene deletion mutants of Δ*fpr3*, Δ*fpr4*, and Δ*sir2* used for the RNA sequencing are all isogenic to BY4741 and were either purchased from open biosystems, or taken from the yeast deletion collection (purchased from Thermofisher Dharmacon). The isogenic double deletion Δ*fpr3*Δ*fpr4* mutant was constructed from the open biosystems Δ*fpr3* single gene deletion mutant by replacing the endogenous *FPR4* locus with a nourseothricin resistance (*MX4-NATR*) PCR product deletion module. The *FPR4*(Δ*fpr3*Δ*trf5*) and Δ*fpr3*Δ*fpr4*Δ*trf5* isogenic strains and their corresponding total haploid mixed population controls were generated from the SGA cross (see below).

The Δ*fpr3* and Δ*fpr4* deletion mutant strains used in the rDNA reporter spotting assays were generated from a cross of the MAT α UCC1188 (Van Leeuwen *et al*, 2002) with a MATa BY4741 deletion mutant see Appendix table 5 for details. The UCC1188 background Δ*fpr3*Δ*fpr4* double deletion mutant, UCC1188 background Δ*sir2* deletion mutant, and HMLα reporter expression mutants were generated by lithium acetate transformation of either UCC1188 or UCC7266 (Van Leeuwen *et al*, 2002) with PCR product deletion modules. The Δ*fpr3*Δ*fpr4* and Δ*sir2* deletion mutant strains used in the propagation assays were generated from a transformation of UCC1188 with PCR product deletion modules.

### Synthetic Genetic Array (SGA) Analysis

SGA analysis was performed using a Singer Instruments ROTOR microbial arraying robot as previously described (Tong & Boone, 2006) with the following modifications. The MAT a/α diploid zygotes resulting from the query strain DMA cross were pinned onto diploid selective YPD + G418/clonNAT plates a total of two times for greater selection against any residual haploids. Sporulation was carried out at room temperature for 14 days. Spores were pinned onto Mat a selective germination media for two rounds of selection as previously described (Tong & Boone, 2006).

The resulting MAT a progeny were subsequently replica plated onto four kinds of selective media: control media selective for the total haploid meiotic progeny population (SD media lacking histidine, arginine, lysine and containing canavanine and thialysine both at a final concentration of 50mg/l, and G418 at a final concentration of 200mg/L), media selective for Δ*xxx*Δ*fpr3* haploid meiotic progeny (SD media lacking histidine, arginine, lysine, leucine, uracil, and containing canavanine and thialysine both at a final concentration of 50mg/l, G418 and clonNAT both at a final concentration of 200mg/L), media selective for Δ*xxx*Δ*fpr4* haploid meiotic progeny (SD media lacking histidine, arginine, lysine, and containing canavanine and thialysine both at a final concentration of 50mg/l, G418 and clonNAT both at a final concentration of 200mg/L, and 5-fluoroorotic acid at a final concentration of 1000mg/L), and finally media selective for Δ*xxx*Δ*fpr3*Δ*fpr4* haploid meiotic progeny (SD media lacking histidine, arginine, lysine, leucine, and containing canavanine and thialysine both at a final concentration of 50mg/l, G418 and clonNAT both at a final concentration of 200mg/L, and 5-fluoroorotic acid at a final concentration of 1000mg/L). Plates were incubated at 30°C for 24 hours and were then expanded into triplicate and incubated for an additional 24 hours at 30°C.

Images of each plate were scanned and subsequently processed using the Balony image analysis software package as previously described (Young & Loewen, 2013). In brief, pixel area occupied by each colony was measured to determine colony size. Progeny fitness was then scored as follows. The ratio of each double (Δ*xxx*Δ*fpr3*, Δ*xxx*Δ*fpr4*) and triple (Δ*xxx*Δ*fpr3*Δ*fpr4*) mutant colony size relative to its corresponding total haploid meiotic progeny control colony was determined. Ratio cutoff thresholds were estimated automatically by the software by extrapolating the central linear portion of the ratio distributions and finding the *y*-intercepts at either ends of the *x*-axis. Default ratio cutoff thresholds were used (a complete list of all genetic interactions generated from each dataset is presented in Appendix file 1).

### SGA Data Processing

Specific, common and masked synthetic sick/lethal interactors were identified as follows. First, duplicate genes from the list of hits from each dataset were identified and removed. The synthetic sick/lethal hits from each of the three datasets were then compared to each other in order to identify unique and common genes in each list. We thus identified a list of interactors unique to the *xxx fpr3* meiotic progeny and a list of interactors unique the *xxx fpr4* meiotic progeny. Hits present in both the *xxx fpr3* meiotic progeny and the *xxx fpr4* meiotic progeny were identified as common interactors of *FPR3* and *FPR4*. Hits that only appear in the *xxxfpr3fpr4* meiotic progeny were identified as masked genetic interactors. Unique, common and masked suppressor interactors were identified the same way.

The lists specific, common, and masked synthetic sick/lethal and suppressor genetic interactors were subsequently analyzed using the web based FunSpec bioinformatics tool (http://funspec.med.utoronto.ca/, Dec 2017). The analysis was performed using a p-value cutoff score of 0.01, and without Bonferroni-correction. A full list of the ontologies uncovered and their corresponding p values is presented in Appendix file 2. Networks illustrating the specific and common genetic interactions were drawn using the Cytoscape software platform (http://www.cytoscape.org/).

### Growth Curves

Growth curves to validate the synthetic sickness phenotypes were carried out as follows. Colonies generated from the SGA assay corresponding to each triple mutant of interest and its respective control colony were isolated and validated for correct genotype by PCR. Confirmed strain isolates were then resuspended in fresh YPD media, normalized to an OD600 of 0.2 and distributed into triplicate wells of a 24 well cell culture plate. Plates were subsequently grown for 16h at 30°C in a shaking plate reader. Readings of OD600 were taken every 30 minutes.

### RNA-Seq Library Preparation and Sequencing

Single colony isolates of each strain were grown to mid log phase in 50ml of liquid yeast extract-peptone-dextrose (YPD) media. Samples were then pelleted and washed once with sterile water before being flash frozen in liquid nitrogen and stored for 16 hours at −80°C. Samples were thawed on ice, and RNA was extracted using a phenol freeze based approach as previously described (Schmitt *et al*, 1990). The extracted RNA was subsequently treated with RNase-free DNase I (Thermo Fisher Scientific) RNA samples were processed and sequenced at the BC Cancer agency Michael Smith Genome Sciences Centre following standard operating protocols. Briefly, total RNA samples were ribo-depleated using the Ribo-Zero Gold rRNA Removal Kit (Yeast) (Illumina) and analyzed on an Agilent 2100 Bioanalyzer using Agilent 6000 RNA Nano Kit (Agilent Technologies, Santa Clara, California). cDNA was generated using the Superscript Double-Stranded cDNA Synthesis kit (ThermoFisher), 100bp paired-end libraries prepared using the Paired-End Sample Prep Kit (Illumina, San Diego, California).

### Processing of Sequencing Data

Sequenced paired-end reads were aligned to the sacCer3 reference genome (https://www.ncbi.nlm.nih.gov/assembly/GCF_000146045.2/) using the BWA aligner (Li & Durbin, 2010) (version 0.6.1-r104-tpx). We observed that out of 5110 *Saccharomyces cerevisiae* genes annotated in Ensembl v90 only 267 are spliced with and most of spliced genes (251) having one intron. Therefore, we considered genomic alignment of RNA-seq reads as a good approximation for the yeast transcriptome analysis. For every library total of ~1.5-2M reads were sequenced, of which ~75-95% of reads were aligned.

To quantify gene expression, we filtered reads that aligned to multiple locations (and therefore can’t be placed unambiguously) by applying a BWA mapping quality threshold of 5. We further collapsed fragments that were duplicated (only counting a single copy of a read pair if a number of pairs with the same coordinates was sequenced) as well removed chastity failed reads and considered only reads that were properly paired. Post-processing was performed using the ‘pysam’ application for python (https://github.com/pysam-developers/pysam). The alignment statistics were calculated using the ‘sambamba’ tool v 0.5.5 5 (Tarasov *et al*, 2015).

We considered cDNA fragment lengths distributions as well as genome-wide distributions of read coverage (data not shown) in order to ensure that these characteristics are similar for the pairs of data sets in the differential gene expression (DE) analysis. Genome wide pair-ended fragment coverage profiles for both strands were generated as well as read counts for every gene for further DE analysis.

The reads-per-kilobase-per-million (RPKM) values were calculated for every gene, and DE analysis was performed using the DEfine algorithm (M.Bilenky et al., unpublished). First, the chi2 p-value was estimated for every gene under the null hypothesis that the gene is not differentially expressed between two data sets. The Benjamini-Hochberg FDR-control procedure was applied (FDR=0.05) to find a p-value threshold. To further reduce noise, we only considered genes with the fold-change (FC) between RPKM values FC>1.5, as well required minimal number of aligned reads >5 per gene. Only reads aligned to the proper strand were considered in the DE analysis.

In addition to the standard DE analysis, where gene expression quantification was done by counting reads falling into the gene boundaries, we considered a model independent approach by calculating read counts in every 175bp long bin genome wide (for both strands), and performed DE analysis between bins (with the same approach we used for genes, see above). After defining the DE bins we overlapped their locations with gene coordinates to determine DE genes. This second approach also provided a list of potential DE expressed intergenic regions. A full list of the DE genes is presented in Appendix file 3.

### Ontology analysis of DE genes

Ontologies associated with differentially expressed genes were identified using the web based FunSpec bioinformatics tool (http://funspec.med.utoronto.ca/, Dec 2018). The analysis was performed on genes displaying a fold change or 1.3 and up using a p-value cut-off score of 0.001, and with Bonferroni-correction. A full list of the ontologies uncovered and their corresponding p values is presented in Appendix file 4.

### Averaged gene read maps

Universal gene coverage profiles were generated as follows; we first crated cDNA fragment coverage profiles genome wide for both strands using all aligned read-pairs. Next, we selected profiles for individual genes and scaled them to 100 units and normalized by the total gene coverage. After that we agglomerated all scaled and normalized gene coverage profiles together. When doing this, the profiles for genes on the negative strand were inverted (in other words we always agglomerate profiles from 5’ to 3’ of gene).

### Spotting assays

The URA3 reporter expression spotting assays were performed in two biological replicates as follows. Freshly grown single colony isolates of each strain were grown in liquid YPD media to mid log phase Cells were subsequently collected, re-suspended in sterile water, and normalized to an OD600=1 (approximately 3×10^7^ cells/ml). The normalized cell suspensions were subjected to 10-fold serial dilutions and 4μl of each dilution was spotted onto standard SD-complete media, SD media without uracil, and SD media with 5-FOA at a final concentration of 1000mg/L and uracil at a final concentration of 50mg/L. Plates were incubated at 30°C and growth was analyzed after 48 hours.

### rDNA Reporter Propagation Assays

The URA+ status of each reporter containing strain was first confirmed by growth on SD media lacking uracil. Saturated overnights were then prepared from single colony isolates of each confirmed strain in liquid YPD media. Cultures were prepared from the overnights in 50ml YPD media and grown at 30°C to mid log phase. Cells were subsequently collected, washed once, resuspended in sterile deionized water, and normalized to an OD600=0.5. Normalized cell suspensions were subsequently diluted 10-fold and 250μl of each dilution was plated on 25ml SD 5-FoA plates. Plates were incubated at 30°C for 16 hours. A total of 96 well-isolated colonies were randomly picked from each 5-FoA plate using the Genetix QPix-2 colony picking robot and deposited onto non-selective solid YPD plates. Plates were incubated for 5 days at 30°C. All 96 colonies on each YPD plate were then replica plated onto SD complete control media and SD media lacking uracil and incubated for 5 days at 30°C before being imaged.

